# A High-Performance Genetically Encoded Fluorescent Biosensor for Imaging Physiological Peroxynitrite

**DOI:** 10.1101/2020.08.17.254771

**Authors:** Zhijie Chen, Shen Zhang, Xinyu Li, Hui-wang Ai

**Author notes:** These authors contributed equally (Zhijie Chen, Shen Zhang, Hui-wang Ai). Correspondence (H.A.).

## Abstract

Peroxynitrite is a highly reactive nitrogen species (RNS) that plays critical roles in signal transduction, stress response, and numerous human diseases. Advanced molecular tools that permit the selective, sensitive, and non-invasive detection of peroxynitrite is essential for understanding its pathophysiological functions. Here, we present pnGFP-Ultra, a high performance, reaction-based, genetically encodable biosensor for imaging peroxynitrite in live cells. pnGFP-Ultra features a *p*-boronophenylalanine-modified chromophore as the sensing moiety and exhibits a remarkable 123-fold fluorescence turn-on response towards peroxynitrite while displaying virtually no cross-reaction with other reactive oxygen/nitrogen species, including hydrogen peroxide. To facilitate the expression of pnGFP-Ultra in mammalian cells, we engineered a highly efficient noncanonical amino acid (ncAA) expression system that is broadly applicable to the mammalian expression of proteins containing various ncAAs. pnGFP-Ultra robustly detected peroxynitrite production during interferon γ and lipopolysaccharide-induced immune responses in macrophages, and in amyloid β-activated primary glial cells. Thus, pnGFP-Ultra fills an important technical gap and represents an important new addition to the molecular toolbox in probing RNS biology.

**In Brief:** Chen et al. report pnGFP-Ultra, a high-performance fluorescent biosensor for minimally invasive and selective imaging of peroxynitrite production in live cells.

**Highlights:**

- pnGFP-Ultra is a genetically encoded peroxynitrite biosensor with a 123-fold fluorescence turn-on response
- pnGFP-Ultra exhibits high selectivity toward peroxynitrite, with virtually no crossreaction with hydrogen peroxide
- An optimized plasmid-based system increases noncanonical amino acid incorporation in mammalian cells by >10 fold
- pnGFP-Ultra robustly detects peroxynitrite production in macrophages and primary glial cells

## Introduction

Reactive oxygen species (ROS) has been studied intensively for decades, but our understanding of its cousin, reactive nitrogen species (RNS) lagged far behind. Peroxynitrite (ONOO^−^) is a RNS formed from a diffusion-controlled reaction between nitric oxide (NO•) and superoxide anion (O_2_^•−^) (Radi, 2013a), and has been implicated in numerous pathophysiological processes, including cardiac (Mungrue et al., 2002; Ronson et al., 1999), vascular (Förstermann and Münzel, 2006), circulatory (Szabó, 1996), inflammatory (Van Der Veen et al., 1997), diabetic (Zou et al., 2004), and neurodegenerative (Smith et al., 1997; Torreilles et al., 1999) diseases. Although no enzyme dedicated for direct peroxynitrite production has been identified, enzymes responsible for the biogenesis of the parental species of ONOO^−^ are widely known: NO• (Beckman and Koppenol, 1996) and O_2_^•−^ (Fridovich, 1997) can be generated by nitric oxide synthetase (NOS) and NADPH oxidase (NOX) (Brewer et al., 2015), respectively (Figure 1A). In addition, the mitochondrial electron transport chain has been recognized as an important source of O_2_’^−^ (Brand, 2016). These primary species present limited cellular toxicity but can react with each other and/or transition metals to produce highly reactive secondary species (*e.g*. hypochlorous acid, hydroxy radical, peroxynitrite) that could have deleterious consequences. For example, the direct toxicity of NO• is modest but can be greatly enhanced by reacting with O_2_^•−^ to produce ONOO^−^. As such, many nitrosative stresses historically attributed to NO• have been gradually reassigned to ONOO^−^ (Pacher et al., 2007), a far more potent RNS capable of mediating radicals (•OH, CO_3_^•−^ and •NO_2_) generation, DNA damage, lipid peroxidation, protein nitration, and cell death (apoptosis or necrosis) (Szabó et al., 2007). In particular, ONOO^−^-mediated post-translational protein tyrosine nitration plays important roles in cell signal transduction (Liaudet et al., 2009). In macrophages, ONOO^−^ protects cells against invading pathogens by acting as a strong oxidant (Allen et al., 2012). Thus, like many reactive oxygen/nitrogen species (ROS/RNS), peroxynitrite displays a sophisticated signal/stress dichotomy that is inextricably linked to its complex biology (Pacher et al., 2007).

**Figure 1.**
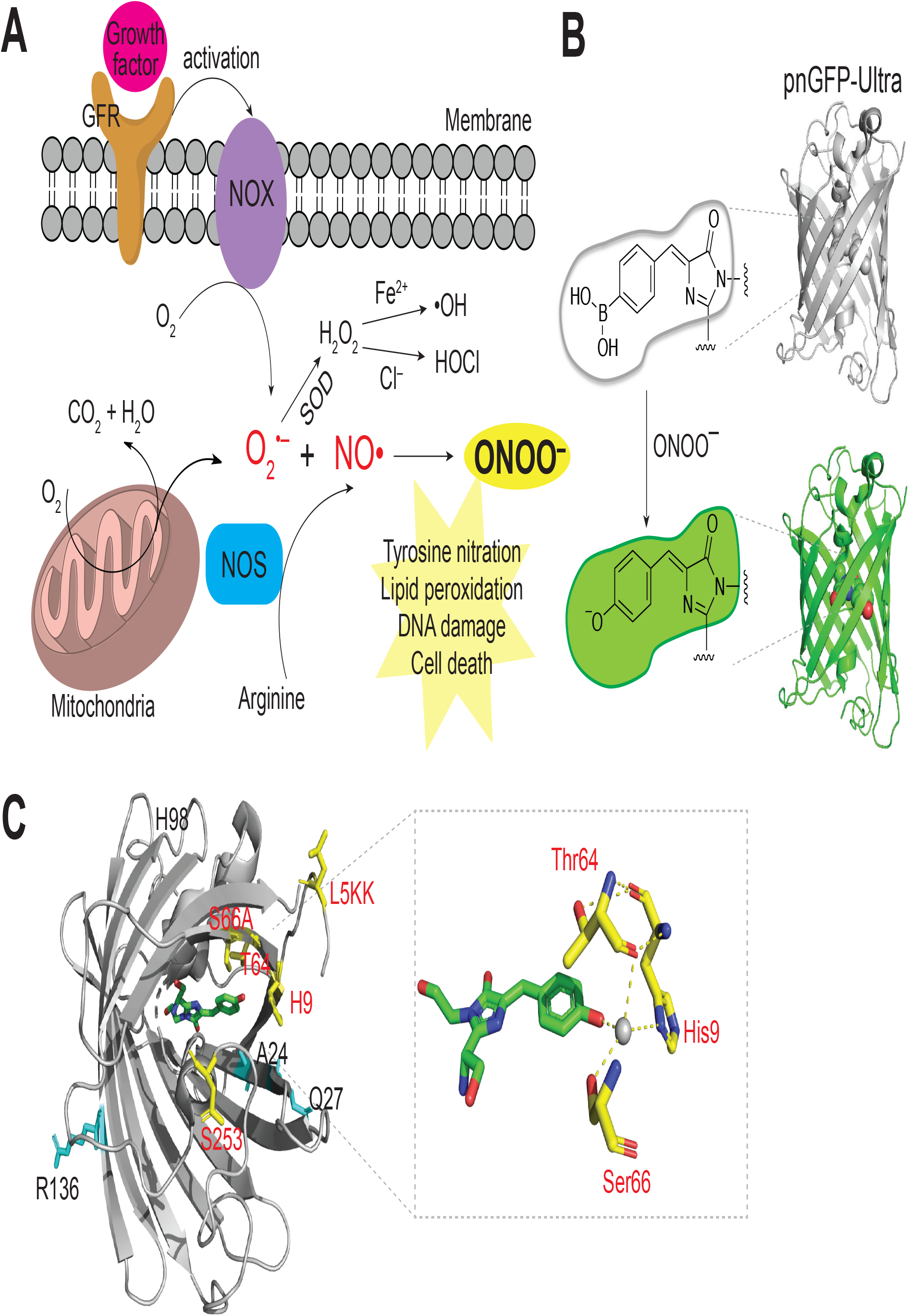
Peroxynitrite Biogenesis and Sensing Mechanism of pnGFP-Ultra. (A) Illustration of the major peroxynitrite (ONOO^−^) biogenesis pathway. ONOO^−^ is generated from the diffusion-controlled reaction between superoxide (O_2_^•−^) and nitric oxide (NOo). O_2_^•−^ could be produced during mitochondrial respiration (oxidative phosphorylation) or by activated NADPH oxidase (NOX). NOX can be activated during cell signaling such as a growth factor binding to a growth factor receptor (GFR). Superoxide dismutase (SOD) can convert O_2_’^−^ to hydrogen peroxide (H_2_O_2_), which can mediate secondary radical (*e.g*. •OH or •OC1) production. NO• is produced by nitric oxide synthetase (NOS), which is coupled to arginine metabolism. ONOO^−^ is a highly reactive nitrogen species that plays crucial roles in tyrosine nitration, lipid peroxidation, DNA damage, and cell death. (B) Sensing mechanism of pnGFP-Ultra. pnGFP-Ultra has a boronic acid-modified chromophore that can be converted to a phenolate chromophore upon reaction with ONOO^−^, resulting in a large fluorescence turn-on response. (C) Cartoon representation of pnGFP1.5-Y.Cro (PDB 5F9G) and key residues targeted during pnGFP-Ultra development. Residues identified during directed evolution that increase the brightness and folding of cpsGFP are highlighted as cyan sticks. Residues that tune of the chemoselectivity of pnGFP-Ultra are highlighted as yellow sticks. The chromophore is show in green. The inset box on the right shows the chromophore and its interacting residues. The grey sphere denotes a water molecule. The structure was prepared using Pymol.

Elucidating the various roles ONOO^−^ plays, whether in the context of normal cell signaling or pathogenesis, is pivotal for developing targeted therapeutics (Beckman, 2009; Szabó et al., 2007). Efforts to study ONOO^−^ encounter similar hurdles to that of studying other ROS/RNS, such as a very short half-life, permeability through and reactions within membrane, multiplex reaction pathways, and signal/stress dichotomy (Ferrer-Sueta and Radi, 2009; Hardy et al., 2018). Traditionally, detection of ONOO^−^ relied primarily on immunostaining of 3-nitrotyrosine footprints, but this method suffers from poor antibody specificity, a requisite for cell lysis, and interference from other nitration sources (Radi, 2013b). More recently, fluorescent probes for ONOO^−^ (Chen et al., 2013; Li et al., 2015; Lin et al., 2013; Peng and Yang, 2010; Peng et al., 2014; Sikora et al., 2009; Sun et al., 2014, 2015, 2009; Tian et al., 2011; Ueno et al., 2006; Xu et al., 2011; Yang et al., 2006; Yu et al., 2011, 2013; Zhang et al., 2012; Zielonka et al., 2010) have emerged as powerful tools for ONOO^−^ detection, due to their high sensitivity, spatial-temporal resolution, and compatibility with well-established fluorescence microscopy platforms (Nadler and Schultz, 2013; Ueno and Nagano, 2011). Deployment of these tools in pathophysiological relevant settings such as in activated macrophage cells (Weber et al., 2020), in live smooth muscles from a mouse model of atherosclerosis (Peng et al., 2014), and in the cerebral vasculature of live mice (Li et al., 2015), has revealed unprecedentedly rich information on peroxynitrite production and pathogenesis. Despite the progress, developing probes with both high reactivity and high selectivity for ONOO^−^ remains a fundamental challenge (Chen et al., 2016a; Hardy et al., 2018). Of all the peroxynitrite probes reported thus far, none have shown absolute selectivity towards ONOO^−^ over other ROS/RNS such as H_2_O_2_, •OH and C1O^−^. For example, HKGreen-4, one of the most advanced probe for ONOO^−^, shows reactivity toward •OH and C1O^−^ at 1 eq probe concentration (1 μM) (Peng et al., 2014); NP3, a probe with high ONOO^−^ sensitivity cross reacts (albeit mildly) with H_2_O_2_ and other species at 2 eq probe concentration (10 μM) (Li et al., 2015); Probe 1–D-fructose shows substantial cross-reaction with C1O^−^ (Sun et al., 2014). In particular, H_2_O_2_, a common cellular ROS that has a long *in vivo* half-life and can be generated at markedly high concentrations (up to low mM) in certain pathophysiological conditions (Brewer et al., 2015), represents a major species competing for probe reactivity in a complex cellular milieu, confounding the unambiguous detection of ONOO^−^. Finally, existing probes—predominately in the form of small molecule dye derivatives—often require organic synthesis and lack the capability to be genetically encoded, a feature that would greatly lower the barrier to the widespread adoption of these probes by other laboratories. A genetically encodable sensor (Chen et al., 2017) would also facilitate cell-/tissue-specific expression for *in vivo* studies, as exemplified by the popular genetically encoded calcium sensors (*e.g*. GCaMP series (Lin and Schnitzer, 2016)).

We previously developed pnGFP (Chen et al., 2013), the first and only genetically encoded fluorescent probe for the selective detection of peroxynitrite, by site-specific incorporation of *p*-boronophenylalanine (*p*BoF) into the chromophore of a circularly permuted superfolder green fluorescent protein (cpsGFP). This is achieved via genetic code expansion (Chin, 2017; Liu and Schultz, 2010), an evolutionary technique that uses an orthogonal aminoacyl-tRNA synthetase (aaRS)–tRNA pair to direct the incorporation of a ncAA into proteins in response to a nonsense codon (often the amber UAG codon). Interestingly, we serendipitously discovered that the boronic acid-based chromophore of pnGFP shows unusual high selectivity for ONOO^−^—essentially nonresponsive to even 1 mM H_2_O_2_ (> 2000 eq probe concentration). Mechanistic studies using site-directed mutagenesis, X-ray crystallography, ^11^B NMR, and computational simulation collectively revealed that the boron atom in pnGFP is converted to an *sp*^3^-hybridized form by a nearby histidine residue via a polarized water bridge (Chen et al., 2016b). The unique pathway by which pnGFP gains high selectivity suggests that the protein scaffold of genetically encoded probes can be engineered to tune their chemoselectivity, which remains challenging to obtain from small molecule probes.

Here, using directed evolution, rational mutagenesis, and targeted reactivity screening guided by the crystal structure of pnGFP1.5-Y.Cro (Chen et al., 2016b), we engineered a high-performance peroxynitrite probe, pnGFP-Ultra, which shows ultra-high sensitivity and selectivity for ONOO^−^. Compared to pnGFP, pnGFP-Ultra has better chromophore maturation, brighter fluorescence, and a 6-fold enhancement on both sensitivity and selectivity, making it the most advanced genetically encoded biosensor for peroxynitrite. To facilitate the use of pnGFP-Ultra in mammalian cells, we systematically optimized the plasmid-based expression system for ncAA incorporation, leading to a 13.3-fold higher *p*BoF-incorporated protein expression in HEK 293T cells. We demonstrate that pnGFP-Ultra can be used to robustly detect physiological peroxynitrite production in activated macrophages and primary mouse glia. Together, these developments make pnGFP-Ultra a powerful tool for imaging peroxynitrite, especially in experiments where genetic encodability and high selectivity are of primary interests.

## Results

### Design, Engineering, and Characterization of pnGFP1.5

pnGFP was engineered by replacing the chromophore tyrosine residue of a circularly permutated superfolder green fluorescent protein (cpsGFP) with a ncAA, *p*BoF (Chen et al., 2013). Upon reaction with ONOO^−^, the boronic acid-based chromophore (dark) is converted into its phenolate form (bright), leading to potent fluorescence turn-on response (Figure 1B). Despite the conceptual success, the broad utility of pnGFP is stymied by several of its drawbacks, including poor protein folding/chromophore maturation, low mammalian cell expression, and dim fluorescence before and after ONOO^−^ conversion. Given the sensing mechanism of pnGFP, we aimed to first improve the folding, expression, and brightness of cpsGFP (Chen and Ai, 2014), the parental template of pnGFP that has a tyrosine derived chromophore (Figure S1). We carried out multiple rounds of directed protein evolution followed by colony-based screening in *Escherichia coli* (*E. coli*) cells (Figures S1A), leading to a brighter mutant, cpsGFP2 (cpsGFP-V24A K27Q Y98H S136R) (Figure S1B). Next, we substituted the chromophore tyrosine residue of cpsGFP2 with *p*BoF (Y174B, where B = *p*BoF) through genetic code expansion. To screen for mutants with high reactivity and selectivity toward ONOO^−^, we performed saturation mutagenesis on threonine 5 (T5) and threonine 253 (T253), two terminal residues that have previously been shown to influence the reactivity of ncAA-containing fluorescent biosensors (Chen and Ai, 2014; Chen et al., 2012, 2013, 2016b). Library screening coupled with fluorescence-based *in vitro* assays (Figure S1) resulted in pnGFP1.5 (cpsGFP2-T5L Y174B T253S) (Figure 2), which exhibited a remarkable 132-fold fluorescence turn-on response towards 100 μM ONOO^−^ (Figure 3A). pnGFP1.5 has a higher protein expression level in *E. coli* as compared to pnGFP, presumably because of enhanced protein folding and chromophore maturation.

**Figure 2.**
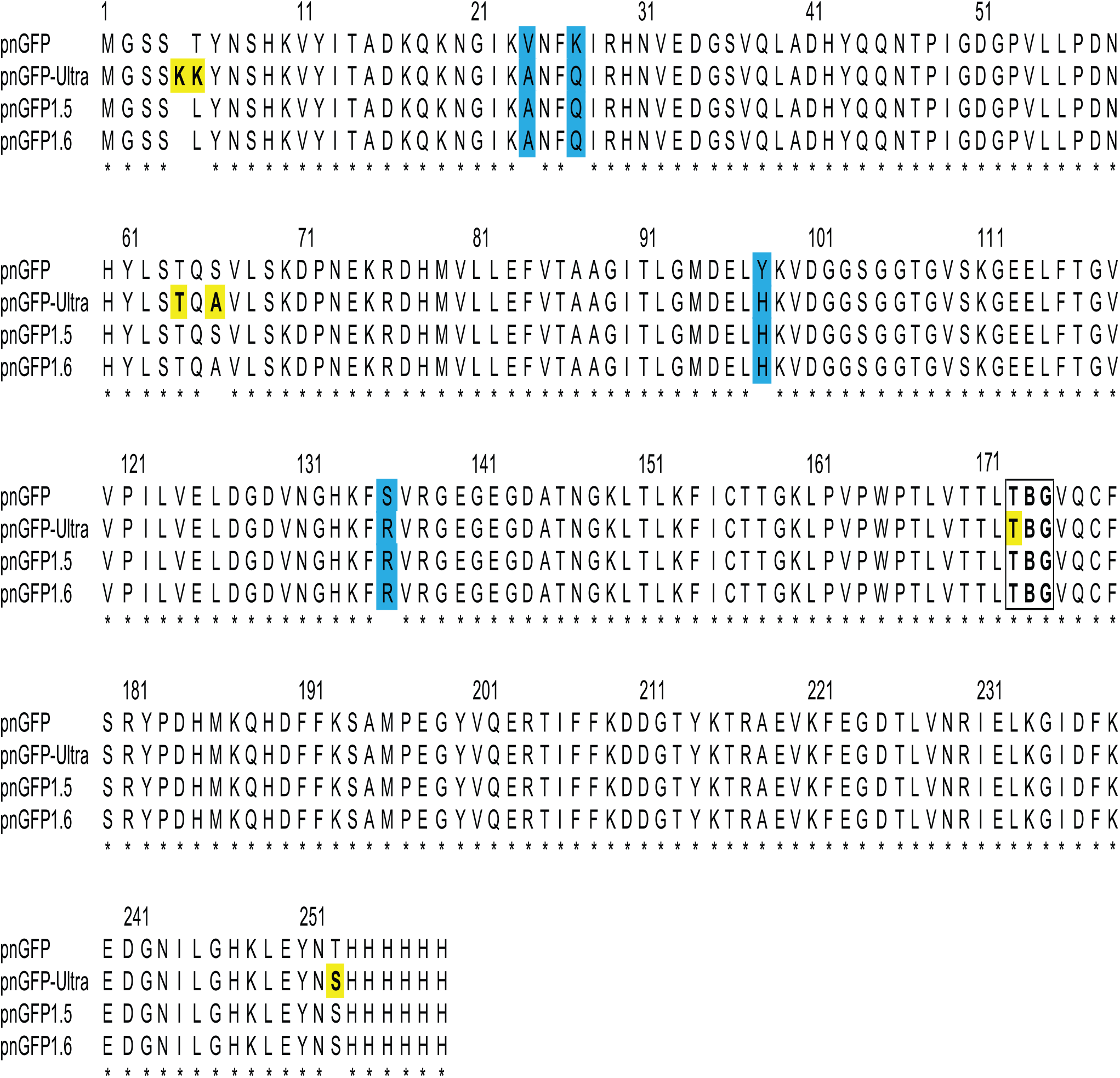
Sequence Alignment of pnGFP-Ultra and Related Proteins. Mutations identified during directed evolution are highlighted in cyan. Residues subjected to rational mutagenesis are yellow-colored. The chromophore-forming residues (B denoting *p*BoF) are highlighted in a box. Residues are numbered according to the numbering of pnGFP1.5-Y.Cro (PDB 5F9G).

**Figure 3.**
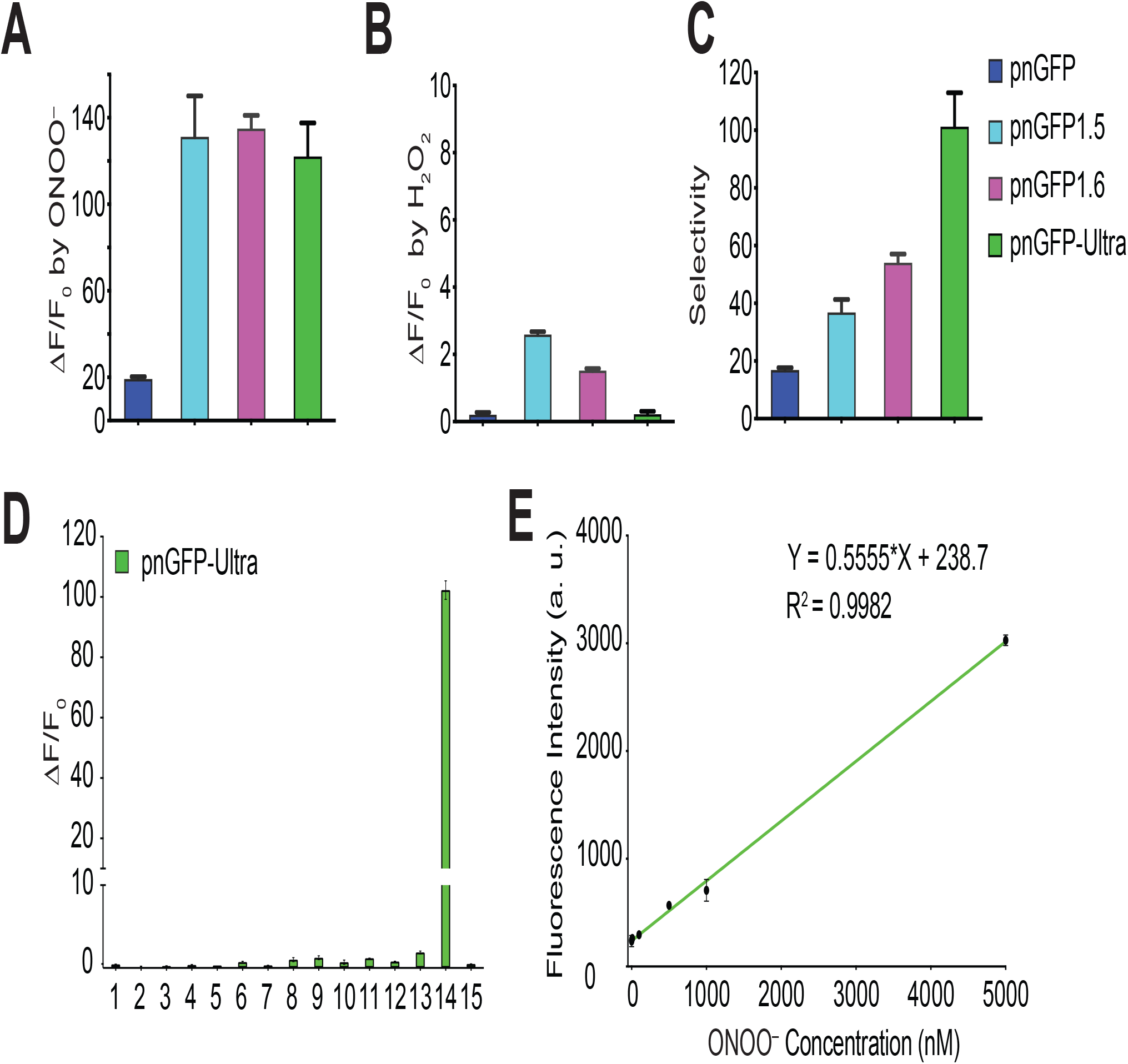
*In vitro* Characterization of pnGFP-Ultra. (A) Dynamic range (expressed as ΔF/F_0_) of pnGFP-Ultra and relevant mutants towards ONOO^−^. F_0_ is the initial fluorescence intensity. ΔF is the final fluorescent intensity (after treatment) minus F_0_. Samples were treated with 100 μM ONOO^−^ for 1 hour. (B) Dynamic range (ΔF/F_0_) of pnGFP-Ultra and relevant mutants towards H_2_O_2_. Samples were treated with 100 μM H_2_O_2_ for 1 hour. (C) Selectivity of pnGFP-Ultra and its mutants. Arbitrarily defined selectivity is calculated as the fold of fluorescence enhancement by ONOO^−^ / fold of fluorescence enhancement by H_2_O_2_. (D) Response of pnGFP-Ultra towards a panel of redox-active chemicals: (1) • OtBu (1 mM Fe^2+^ and 100 μM HOOtBu), (2) 100 μM HOOtBu, (3) • OH (1 mM Fe^2+^ and 100 μM H2O2), (4) 5 mM oxidized glutathione, (5) 100 μM O_2_^•−^, (6) 100 μM NOC-7 (NO• donor), (7) 100 μM HOC1, (8) 100 μM NaHS (H_2_S donor), (9) 5 mM L-cysteine, (10) 1 mM DL-homocysteine, (11) 1 mM vitamin C, (12) 100 μM H2O2, (13) 1 mM H_2_O_2_, (14) 100 μM ONOO^−^, (15) Tris buffer. (E) Linear fluorescence responses of pnGFP-Ultra (0.2 μM) to ONOO^−^ from high nanomolar to low micromolar concentrations. The limit of detection was determined to be 277 nM with a signal-to-noise ratio (S/N) of 3.

### Structure-Guided Engineering of pnGFP-Ultra

Compared to pnGFP, pnGFP1.5 showed a 6-fold enhancement in the dynamic range towards ONOO^−^ (Figure 3A), but also an undesirable ~3-fold increase in the dynamic range towards H_2_O_2_ (Figure 3B). The overall selectivity (arbitrarily defined as fold of fluorescence enhancement by ONOO^−^ / fold of fluorescence enhancement by H_2_O_2_) of pnGFP1.5 remains higher than that of pnGFP (Figure 3C). We next sought to tune the selectivity of pnGFP1.5 by rational mutagenesis. In our previous attempt to decipher the unusual chemoselectivity of pnGFP, we solved the X-ray crystal structure of pnGFP1.5-Y.Cro (pnGFP1.5-B174Y, PDB 5F9G), which contains a tyrosine-derived chromophore (Chen et al., 2016b). The structure indicates that three residues—namely Thr64, Ser66, His9—are in close proximity to the phenolate oxygen of the chromophore (Figure 1C). We speculate that these residues may interact with the *p*BoF-derived chromophore in pnGFP1.5 to fine tune the chromophore environment (electrostatics, hydrophobicity, and solvent accessibility). In addition, the first chromophore-forming residue (Thr173) can also affect electron distributions within the conjugate chromophore structure. Thus, these four residues are next subjected to site-directed mutagenesis (Figure 2 and S1). Based on our prior experiences with pnGFP, we created a panel of pnGFP1.5 mutants and assayed their selectivity *in vitro* (Figure S2). From this screening, we identified pnGFP1.6 (pnGFP1.5-S66A), which showed much lower reactivity towards H_2_O_2_, but maintained the same amplitude of response towards ONOO^−^. All other mutants exhibited lower selectivity than pnGFP1.5, either due to low fluorescence intensity after ONOO^−^ conversion or high reactivity towards H_2_O_2_ (Figure S2). We next focused on improving pnGFP1.6 (Figure S1B).

pnGFP gains selectivity, in part, by leveraging the nucleophilic attack efficiency differences between ONOO^−^ and HOO^−^ (Chen et al., 2016b). Due to *p*Ka differences (11.6 for H_2_O_2_ and 6.8 for ONOOH), the portion of HOO^−^ at neutral pH is much lower than that of ONOO^−^, thereby disfavoring the reaction of pnGFP with H_2_O_2_. However, this difference alone is not sufficient to explain the remarkable selectivity of pnGFP, suggesting that the protein scaffold contributes to further reducing the effective local concentration of HOO^−^ available for reaction with the *p*BoF-derived chromophore. To this end, we substituted threonine 5—which gates an opening towards the chromophore—with two randomized residues by using degenerate NNK codons. Screening of the resultant library against both H_2_O_2_ and ONOO^−^ led to pnGFP-Ultra (pnGFP1.6-T5KK, Figure 2 and S1B), which showed virtually no response to 100 μM H_2_O_2_ even after 1-hour incubation (Figure 3B). While this came at a price of a slightly reduced reactivity to ONOO^−^ (Figure 3A), the selectivity of pnGFP-Ultra is 1.9- and 6-fold that of pnGFP1.6 and pnGFP, respectively (Figure 3C). Interestingly, in pnGFP-Ultra, threonine 5 is replaced with two consecutive lysine residues (Figure 2), which possibly abolish the reactivity of pnGFP-Ultra towards H_2_O_2_ by channeling a positively charged entry route that diminishes local availability of H_2_O_2_ near the *p*BoF-derived chromophore. This unexpected screening result further highlights the unique scaffolding effect of protein-based sensors, which could be harnessed to achieve unusual chemoselectivity.

We tested *in vitro*-purified pnGFP-Ultra against a panel of redox-active molecules, including ROS/RNS and thiols that are commonly present in a complex cellular milieu (Figure 3D). Except for ONOO^−^, none of the species surveyed triggered appreciable fluorescent changes, confirming that pnGFP-Ultra is highly selective towards ONOO^−^ (Figure 3D). Immediately (less than 10 seconds) upon incubation with 100 μM ONOO^−^, pnGFP-Ultra displayed a robust 123-fold fluorescence enhancement. By contrast, 1-hour incubation of pnGFP-Ultra with 100 μM and 1 mM H_2_O_2_ triggered only 1.31- and 2.46-fold fluorescence increase, respectively (Figure 3D). Given the low basal fluorescence intensity of pnGFP-Ultra (essentially dark), these small fluorescence increases are negligible in most applications. The fluorescence intensity of pnGFP-Ultra increased linearly from high nanomolar to low micromolar ONOO^−^ concentrations, with the limit of detection determined to be 277 nM (Figure 3E). Compared to pnGFP, pnGFP-Ultra exhibits an overall 6-fold improvement in both the dynamic range and selectivity (Figure 3A and 3C). Taken together, these results demonstrate that pnGFP-Ultra is a high performance RNS biosensor that is highly sensitive and selective for peroxynitrite.

### Development of an Enhanced ncAA Incorporation System in Mammalian Cells

Efficient co-translational ncAA incorporation in response to the amber TAG codon is crucial for the success of pnGFP-Ultra and other applications that require ncAA incorporation. Indeed, one of the fundamental limitations of pnGFP and other ncAA-based biosensors is their inadequate expression in mammalian cells. While ncAA incorporation in *E. coli*. has been thoroughly optimized to an extent that ncAA incorporation can reach a level comparable to that of canonical amino acids, mammalian incorporation of ncAA has proven to be far more challenging, presumably due to a more complicated ribosomal translation machinery.

Existing plasmid-based systems for ncAA incorporation (Chen et al., 2009; Hino et al., 2012; Liu et al., 2007; Schmied et al., 2014; Xiao et al., 2013) vary in the type and copy number of the engineered orthogonal tRNA-aminoacyl tRNA synthetase pair, and the choices of vectors and promoters, making it difficult to cross compare the efficiencies of these systems. Several strategies for enhancing amber codon suppression efficiency such as increasing the copy number of tRNAs (Liu et al., 2007) and ectopic expression of a dominant-negative eukaryotic release factor 1 mutant (eRF1-E55D) (Schmied et al., 2014) have proven to be generalizable. To systematically optimize ncAA incorporation into proteins in mammalian cells, we constructed a series of expression plasmids (plasmids *a-e*, Figure 4A) and tested ncAA incorporation in HEK293T cells using an EGFP_TAG_ fluorescent reporter gene that bears an amber TAG codon at position 39 of EGFP. Incorporation of a ncAA in response to the amber TAG codon (amber suppression) would generate the full length EGFP while failure to do so would result in a truncated EGFP (non-fluorescent). Previously, *p*BoF was incorporated into pnGFP by co-transfecting the CMV promoter-driven pnGFP reporter plasmid with the suppressor plasmid, which harbors one copy of CMV promoter-driven poly-specific aminoacyl tRNA synthetase (Poly-aaRS) and four copies of U6/H1 promoter-driven suppressor tRNAs (Figure 5A). Since orthogonal tRNA expression is a limiting factor in ncAA incorporation efficiency, we cloned four copies of tRNAs on the reporter plasmid *c* as well (Figure 4A). Compared to our previous expression method using plasmids *a + b*, addition of extra copies of tRNA on the reporter plasmid doubled the incorporation efficiency of *p*BoF (Figure 4B) into the EGFPTAG reporter (Figure 4C, plasmids *c + b*). To test whether ectopic expression of eRF1-E55D further enhances ncAA incorporation efficiency, eRF1-E55D was bicistronically expressed with Poly-aaRS via a self-cleaving T2A peptide linker (Figure 4A, plasmid *d*). Co-expression of eRF1-E55D-containing suppressor plasmid *d*, in lieu of *b*, with the original reporter plasmid *a* resulted in a 10.9-fold enhancement in *p*BoF incorporation (Figure 4C). Remarkably, we observed further improvements in incorporation efficiency using plasmids *c + d*, suggesting that the benefit from expression of extra copies of tRNA and the eRF1-E55D mutant are synergistic (Figure 4C). To simply the expression system, we integrated all genetic components necessary for ncAA incorporation—including the reporter gene, Poly-aaRS, and orthogonal tRNAs—into a single plasmid (Figure 4A, plasmid *e*). Expression of plasmid *e* alone gave rise to comparable incorporation efficiency to that of using *a + d*. Co-transfection of plasmid *e* with other plasmids led to various degrees of expression efficiency. Nevertheless, all strategies tested surpass our original method of using plasmids *a + b* for ncAA incorporation, with plasmids *c + d* and *c + e* having the highest incorporation efficiency for *p*BoF (13.3-fold enhancement) (Figure 4C). We achieved similar enhancement in incorporation efficiency for *p*-azido-phenylalanine (*p*AzF) (Figure S3), suggesting that this plasmid-based system is quite general and can be used to incorporate other ncAAs.

**Figure 4.**
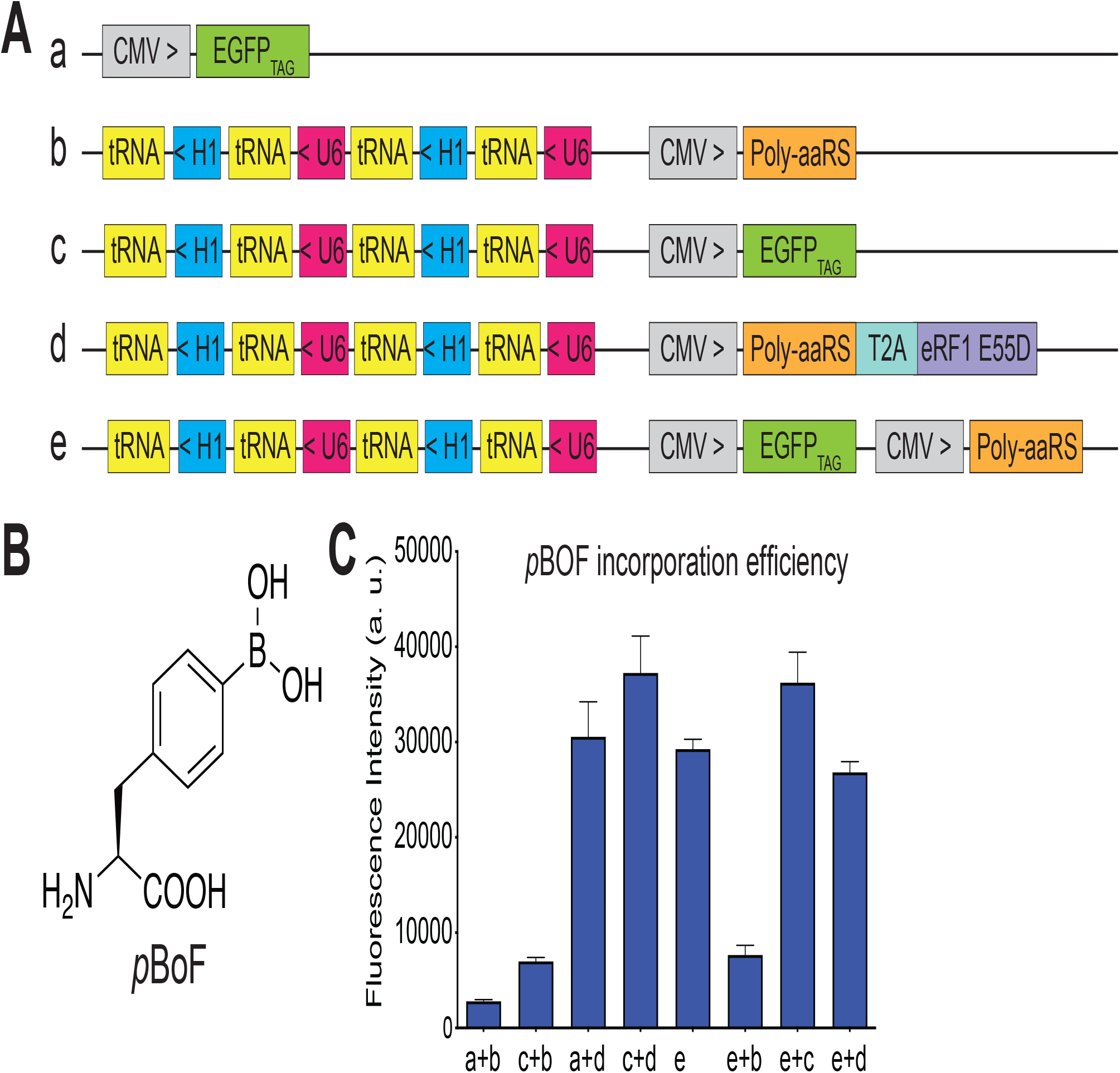
Engineering of a Highly Efficient Plasmid-based ncAAs Incorporation System in Mammalian Cells. (A) Schematics of various expression plasmids. Poly-aaRS is the poly-specific aminoacyl tRNA synthetase, eRF1-E55D is the mutant eRF1 gene, T2A is a self-cleaving 2A peptide, tRNA is the orthogonal tRNA, U6 indicates the U6 promoter, H1 indicates the H1 promoter, CMV is the CMV promoter, EGFP_TAG_ is the EGFP reporter gene with a TAG codon at position 39. > or < denotes the direction of the gene cassettes. (B) Chemical structure of *p*-Boronophenylalanine (*p*BoF). (C) Quantification of *p*BoF (1 mM) incorporation into the EGFP_TAG_ reporter gene measured in a cell-based fluorescence assay. The indicated constructs were transiently expressed in HEK 293T cells and the cell lysate green fluorescence was quantified in a plate reader at 515 nm emission with excitation at 490 nm. Data represent the mean ± SD of triplicates.

**Figure 5.**
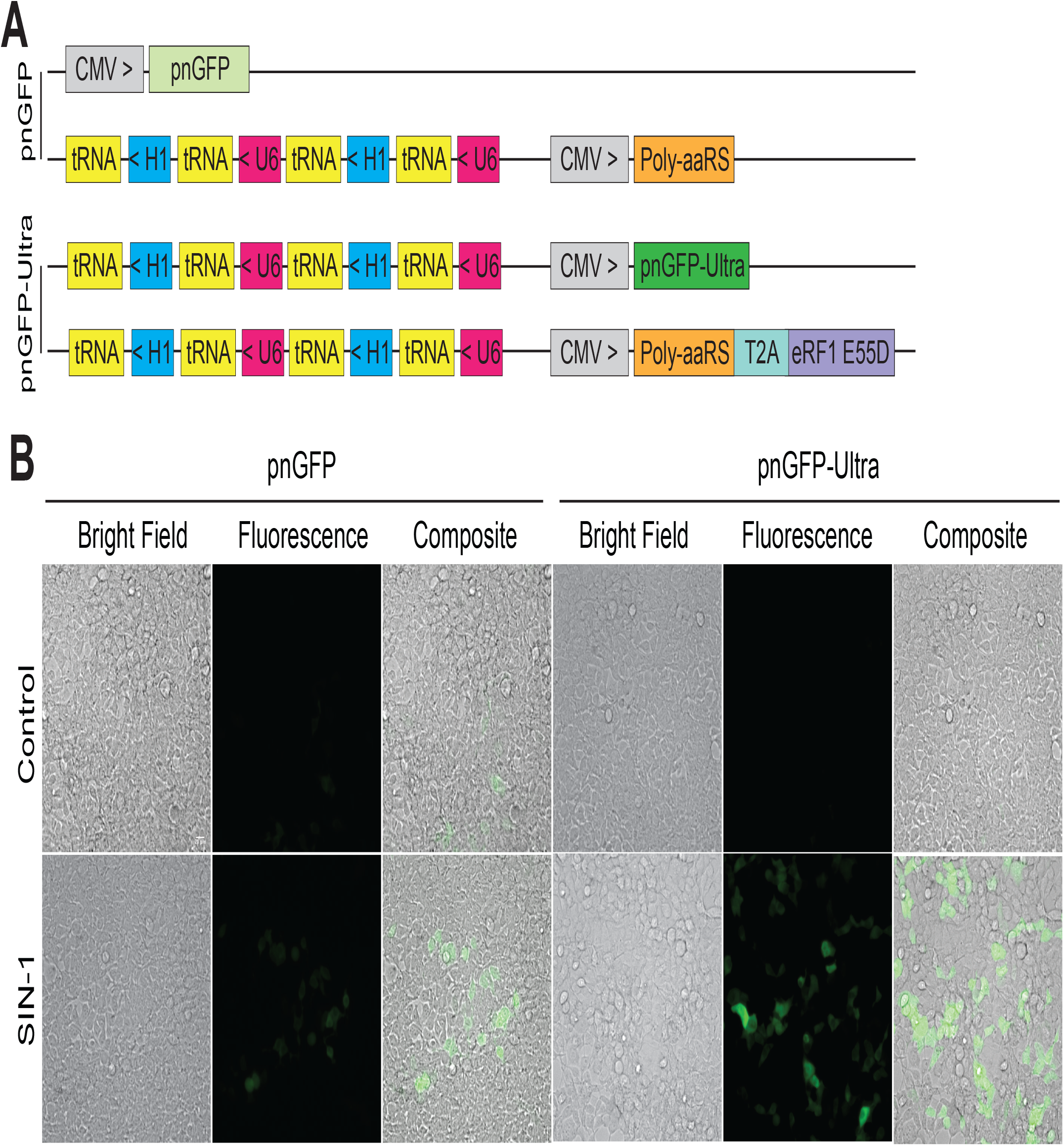
Live-cell imaging of peroxynitrite in HEK 293T cells. (A) Constructs used for expressing pnGFP (top two plasmids) and pnGFP-Ultra (bottom two plasmids). (B) Green fluorescent and bright-field images of cells expressing pnGFP (left) or pnGFP-Ultra (right). In the bottom row, HEK 293T cells were treated with 100 μM SIN-1 for 90 min before imaging. The number of fluorescent cells from the pnGFP-Ultra-transfected group was approximately 2-fold of that from the pnGFP-transfected group.

### Imaging of Peroxynitrite in Live Cells using pnGFP-Ultra

Next, we constructed a mammalian expression plasmid by replacing the EGFP_TAG_ gene with pnGFP-Ultra gene in plasmid *c* (Figure 5A). We co-expressed the resultant plasmid with plasmid *d*, as this combination gave rise to the highest *p*BoF incorporation efficiency in the above EGFP_TAG_ reporter assay (Figure 4C). To compare with the first-generation biosensor, we expressed pnGFP in parallel by utilizing our previous method (Figure 5A). After treatment with SIN-1, a cell permeable ONOO^−^ donor, the fluorescence intensities from pnGFP-Ultra-expressing cells were much higher than those from pnGFP-expressing cells (Figure 5B). It is also evident (from the number of fluorescent cells) that the expression efficiency of pnGFP-Ultra is much higher than that of pnGFP. Both pnGFP and pnGFP-Ultra are selective towards ONOO^−^, with no obvious response toward H_2_O_2_ (1 mM) or HOCl (100 μM) (Figure S4). We attribute the superior performance of pnGFP-Ultra to both its large dynamic range (123-fold fluorescence enhancement) and the substantially improved *p*BoF incorporation. Together, these developments make pnGFP-Ultra a high-performance biosensor for imaging ONOO^−^ in mammalian cells.

### Visualization of Peroxynitrite Production in Activated Macrophages

To further demonstrate the utility of pnGFP-Ultra in imaging physiological peroxynitrite, we expressed pnGFP-Ultra in mouse RAW264.7 macrophage immune cells, which are known to produce ONOO^−^ for microbicidal activities under immune stimulation (Allen et al., 2012; Peng et al., 2014; Xia and Zweier, 1997). Upon co-stimulation with bacterial endotoxin lipopolysaccharide (LPS) and pro-inflammatory cytokine interferon-gamma (IFN-γ), we observed strong fluorescence signal in pnGFP-Ultra expressing cells (Figure 6A). The mean fluorescence of cells in the treated group was ~ 3.7-fold higher than that of untreated cells (Figure 6B), suggesting that pnGFP-Ultra can reliably detecting physiological peroxynitrite production.

**Figure 6.**
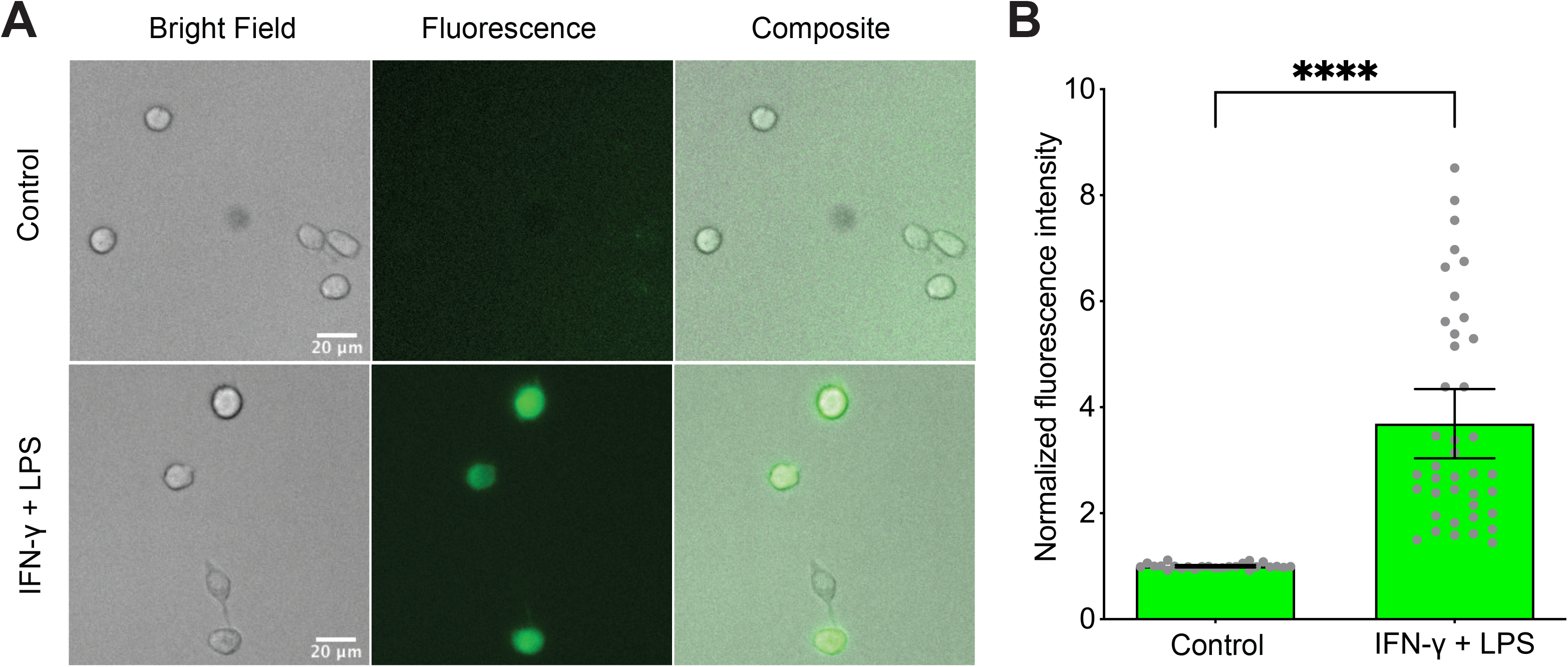
Live-cell Imaging of Peroxynitrite Production in Activated Macrophages. (A) Mouse RAW264.7 cells were transfected with pnGFP-Ultra. After 24 hours, the samples were untreated or treated with IFN-γ (50 U/mL) and LPS (100 ng/mL) for 36 hours before imaging. Representative bright field (left), fluorescence (middle) and overlay (right) images of each group were shown. (B) Quantification of fluorescence intensities of cells untreated or treated with IFN-γ and LPS. Gray dots indicate fluorescent intensities of single cells. Box plots represent Mean ± 95% confidential intervals (n = 25 individual cells for each group). Statistical test was performed using Welch’s unpaired t test (**** indicate P < 0.0001).

### Detection of Peroxynitrite Production in Amyloid β-Activated Primary Mouse Glia

Despite the continuous improvement in life expectancy, Alzheimer’s disease (AD) and other neurodegenerative disorders have become a global health and social challenge (Association, 2019). Peroxynitrite is known to play important roles in numerous neurodegenerative diseases. For example, peroxynitrite-mediated damage has been broadly observed in AD (Smith et al., 1997) and the amyloid β peptide, whose accumulation has been considered to be the hallmark of AD, is known to induce peroxynitrite production in glial cells (González-Reyes et al., 2017; Xie et al., 2002) and subsequently mediate neurotoxicity (Xie et al., 2002). To test whether pnGFP-Ultra can detect peroxynitrite production in neurological systems, we expressed pnGFP-Ultra in cultured primary mouse glia cells. Indeed, we observed dramatically enhanced fluorescence signal from glia activated with amyloid β (Figure 7A), producing a ~3.8-fold mean fluorescence enhancement compared with untreated cells (Figure 7B). Thus, pnGFP-Ultra represents a novel tool enabling the imaging of peroxynitrite production in neurological systems. In addition, it opens up an exciting possibility to develop pnGFP-Ultra-based fluorescent assays for high-throughput screening of inhibitors that targets major ONOO^−^ production pathways. Such capability will aid the development of drugs that protect neurons against ONOO^−^-mediated cell toxicity/death (Szabó et al., 2007).

**Figure 7.**
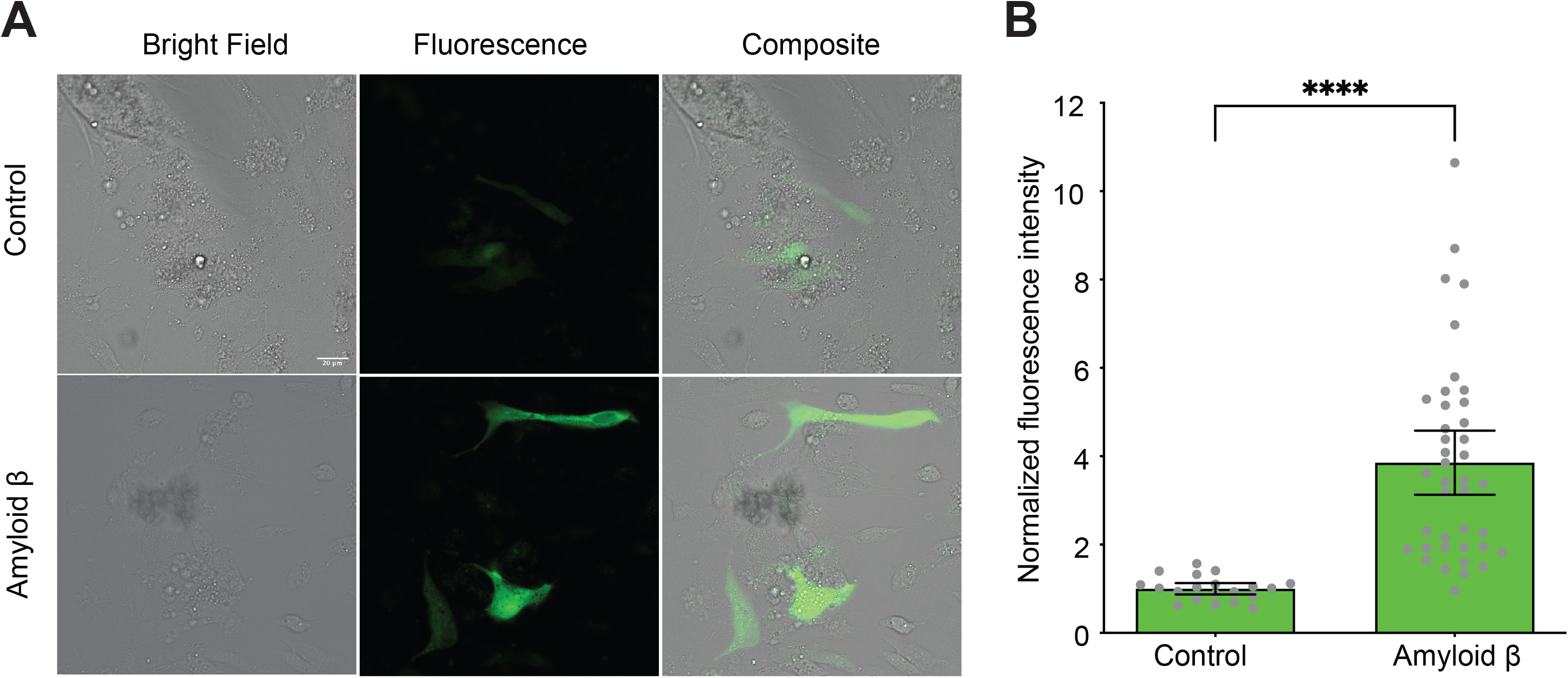
Live-cell Imaging of Amyloid β-Induced Peroxynitrite in Primary Glial Cells. (A) Primary mouse glia cells were isolated and transfected with pnGFP-Ultra. After 24 hours, the samples were untreated or treated with amyloid β (5 μM) for 42 hours before imaging. Representative bright field (left), fluorescence (middle) and overlay (right) images of each group were shown. (B) Quantification of fluorescence intensities of cells untreated or treated with amyloid β. Gray dots indicate fluorescent intensities of single cells. Box plots represent Mean ± 95% confidential intervals (n = 20 and 40 individual cells for control and amyloid β-treated groups, respectively). Statistical test was performed using Welch’s unpaired t test (**** indicate P < 0.0001).

## Discussion

In summary, we have developed pnGFP-Ultra—a highly sensitive and selective fluorescent peroxynitrite biosensor—by directed evolution, structure-guided rational design, and reactivity-based library screening. Compared to the first-generation peroxynitrite probe pnGFP, pnGFP-Ultra has much better chromophore maturation and a 6-fold larger ONOO^−^-induced fluorescence response, while retaining the high chemoselectivity against H_2_O_2_ and other ROS/RNS. An apparent hurdle of applying ncAA-based biosensors to cell imaging studies has been the inefficiency with which ncAA can be incorporated into proteins in mammalian cells. Here, we systematically optimized and increased *p*BoF incorporation efficiency in HEK 293T cells by 13.3-fold. These developments make pnGFP-Ultra a high-performance biosensor capable of imaging physiological peroxynitrite, as we have demonstrated in activated macrophages and primary glial cells. pnGFP-Ultra thus represents an important advancement enabling the practical implementation of a genetically encoded biosensor for live-cell detection of peroxynitrite. We expect pnGFP-Ultra to find broad utility in probing RNS biology in diverse pathophysiological settings.

pnGFP-Ultra not only fills an important technological gap in the field of RNS, but also conveys innovations that will inspire the development of future biosensors for other bioanalytical targets. First, the chromophore (and other residues) of circularly permutated fluorescent proteins can be modified into a hub of chemical sensing, binding, reactivity, and catalysis, thereby greatly expanding the use of fluorescent proteins as reporters. Second, direct evolution and rational mutagenesis performed on noncanonical amino acid-modified proteins can lead to new and sometimes, surprising functions, as we have demonstrated here via the engineering of exquisite chemo-selectivity of pnGFP-Ultra.

Compared to existing small-molecule based peroxynitrite probes, pnGFP-Ultra has the advantage of being genetically encodable. This feature permits long-term *in vivo* imaging and organelle-/cell-/tissue-specific targeting, making it possible to apply pnGFP-Ultra to a variety of biological/disease models. More importantly, because of the convenient fluorescence readout and robust off-on response of pnGFP-Ultra, cell lines stably expressing this reporter can be used in high-throughput drug screening assays. pnGFP-Ultra may also be used in genome-wide CRISPR screens (Shalem et al., 2015) to uncover new genes involved in NO•, O_2_^−•^, or ONOO^−^ production pathways. In addition to live cell imaging, we envision these high-throughput drug/genetic screens to be an important avenue of application where pnGFP-Ultra shall prove to be an enabling tool. Since pnGFP-Ultra can be widely distributed in the form of plasmids, it represents a valuable catalyst to future breakthroughs in the field of RNS.

## STAR⍰Methods

### Key Resources Table

| REAGENT or RESOURCE | SOURCE | IDENTIFIER |
| --- | --- | --- |
| **Chemicals, Peptides, and Recombinant Proteins** |  |  |
| *p*-borono-DL-phenylalanine | Synthonix | Cat#: B5001;<br>CAS:90580-64-6 |
| *p*-azido-phenylalanine | Bachem | Cat#: F-4305;<br>CAS:1241681-80-0 |
| Sodium peroxynitrite | Cayman Chemical | Cat#: 81565;<br>CAS:14042-01-4 |
| 3-morpholinosydnonimine (SIN-1) | Sigma Aldrich | Cat#: M5793;<br>CAS:16142-27-1 |
| Human Interferon-γ Protein, recombinant | Sigma Aldrich | Cat#: IF002 |
| Lipopolysaccharides (LPS)<br>from *Salmonella typhosa* | Sigma Aldrich | Cat#: L7895 |
| β-Amyloid (1-42), human | GenScript | Cat#: RP10017 |
| **Critical Commercial Assays** |  |  |
| B-PER | Thermo Fisher Scientific | Cat#: 78243 |
| TRI Reagent | Thermo Fisher Scientific | Cat#: AM9738 |
| SuperScript® IV Reverse Transcriptase | Thermo Fisher Scientific | Cat#: 18090010 |
| RIPA Lysis and Extraction Buffer | Thermo Fisher Scientific | Cat# 89900 |
| **Cell Culture Reagents** |  |  |
| Dulbecco's modified Eagle's medium (DMEM) | Corning | Cat# 10-017-CV |
| Trypsin EDTA | Corning | Cat# 25-053-CI |
| Fetal Bovine Serum | GenClone | Cat# 25-550 |
| Penicillin-Streptomycin Solution, 100x | Corning | Cat# 30-002-CI |
| β-mercaptoethanol | MP Biomedicals | Cat# 773648<br>CAS: 60-24-2 |
| DPBS | Corning | Cat# 21-030-CVR |
| **Cell Transfection Reagents** |  |  |
| Viomer | ORIGENE | Cat# TT100312 |
| Lipofectamine 2000 | Thermo Fisher Scientific | Cat# 11668027 |
| **Data** |  |  |
| MS source data | This paper | Available upon request |
| **Experimental Models: Cell Lines** |  |  |
| Human HEK 293T cells | ATCC | CRL-3216™ |
| *Escherichia coli* DH10B competent cells | Thermo Fisher Scientific | Cat# 18297010 |
| RAW264.7 Macrophage cells | ATCC | TIB-71™ |
| **Recombinant DNA** |  |  |
| Synthetic oligo nucleotides | IDT | Table S1 |
| pUltra | (Lou et al., 2012) | Addgene # 24129 |
| pMAH-POLY | (Chen et al., 2013) | Addgene # 64915 |
| pcDNA3-pnGFP | (Chen et al., 2013) | Addgene # 64913 |
| pEvol-BoF | (Young et al., 2010) | N/A |
| pBAD-cpsGFP2 | This paper | N/A |
| pBAD-pnGFP-Ultra | This paper | Addgene #157923 |
| pcDNA3-EGFP<sub>TAG</sub> | This paper, Figure 4A, a | N/A |
| pMAH-EGFP<sub>TAG</sub> | This paper, Figure 4A, c | N/A |
| pMAH-POLY-eRF1<sub>E55D</sub> | This paper, Figure 4A, d | Addgene #157925 |
| pMAH-EGFP<sub>TAG</sub>-POLY | This paper, Figure 4A, e | N/A |
| pMAH-pnGFP-Ultra | This paper, Figure 5A | Addgene #157924 |
| **Softwares and Algorithms** |  |  |
| Fiji | (Schindelin et al., 2012) | https://fiji.sc |
| PyMOL | Schrödinger | https://pymol.org/2/ |
| GraphPad Prism 8 | GraphPad | https://www.graphpad.com/scientific-software/prism/ |

### Resource Availability

#### Lead Contact

Further information and requests for resources and reagents should be directed to and will be fulfilled by the Lead Contact, Huiwang Ai.

#### Materials Availability

All reagents created in this study (see Key Resources Table) are available on request. Key plasmids generated have been deposited to Addgene.

#### Data Availability

Raw source data are available upon reasonable request.

#### Experimental Model and Subject Details

HEK 293T cells (human, female, embryonic kidney) were maintained in Dulbecco’s modified Eagle’s medium (DMEM) supplemented with 10% (v/v) FBS at 37°C and 5% CO_2_ atmosphere. Other cell types were cultured as described in Supplementary Experimental Procedures.

## Methods Details

### Materials and General Methods

Synthetic DNA oligonucleotides were purchased from Integrated DNA Technologies (San Diego, CA). Restriction endonucleases were purchased from Thermo Fisher Scientific (Waltham, MA). PCR and restriction digestion products were purified by gel electrophoresis and extracted using the Syd Labs Gel Extraction kit (Malden, MA). Plasmid DNA was purified using the Syd Labs Miniprep kit (Malden, MA). DNA sequencing was analyzed by Retrogen (San Diego, CA). The amino acid *p*-borono-DL-phenylalanine (*p*BoF) was purchased from Synthonix (Wake Forest, NC). Fluorescence measurements were performed on a monochromator-based Synergy Mx Microplate Reader (BioTek, Winooski, VT). Mammalian cell imaging was performed on a Leica SP5 confocal fluorescence microscope or a Leica DMi8 microscope equipped with a Photometrics Prime 95B sCMOS camera. Other materials and general procedures were acquired or performed as previously described (Chen et al., 2012, 2013, 2016b).

### Engineering and Screening of pnGFP1.5

To improve the brightness and folding of pnGFP, several rounds of error prone PCR (EP-PCR) based directed evolution were performed to improve its template—cpsGFP (Chen and Ai, 2014), which contains a tyrosine-derived chromophore. Briefly, oligos pBAD-FP and pBAD-RP were used to amplify the cpsGFP gene from the pBAD-cpsGFP plasmid in an EP-PCR reaction condition. The mutated PCR products were digested with XhoI and HindIII and ligated into a predigested compatible pBAD vector. The ligated product was used to transform *Escherichia coli* DH10B competent cells by electroporation. Cells were grown on Luria-Bertani (LB) broth agar plates supplemented with 100 μg/mL ampicillin and 0.02% L-arabinose. LB agar plates were incubated at 37°C overnight. Bacterial colonies were illuminated under a laboratory-built colony fluorescence imaging system and examined using a pair of forensic yellow goggles with a cutoff wavelength of ~490 nm. Plasmid DNA from the brightest colonies of each round were extracted and combined to serve as the template for the next round of EP-PCR. The fluorescence of the B-PER (Pierce, Rockford, IL) extracted cell lysate from the brightest colonies of the final round was quantitatively compared on the Synergy Mx Microplate Reader. The best mutant was sequenced and renamed as cpsGFP2.

To screen for pnGFP1.5, the codon for the chromophore tyrosine (Tyr174) of cpsGFP2 were mutated to TAG to allow for *p*BoF incorporation using the pEvol-BoF-based amber suppression system. Next, threonine 5 (Thr5) and threonine 253 (Thr253) of cpsGFP2-Tyr174TAG were randomized using NNK degenerate codons to construct a gene library. The library was used to screen for mutants with high reactivity to ONOO^−^. The chromophore TAG mutagenesis, library construction and screening procedures were the same as those in pnGFP engineering, which was detailed previously (Chen et al., 2013). The best mutant from the library was sequenced and designated as pnGFP1.5.

### Development and Characterization of pnGFP-Ultra

To create the various pnGFP1.5 mutants, site-directed mutagenesis of pnGFP1.5 was performed using an overlap extension PCR based strategy. The procedures were described previously (Chen et al., 2016b), and the mutagenesis primers can be found in Supplementary Information Table S1. Proteins from each mutant were purified, buffer-exchanged and concentrated as described. To test the reactivity and selectivity of the pnGFP1.5 mutants, 0.2 μM purified protein was incubated with Tris buffer (150 mM Tris-HCl, 150 mM NaCl, pH 7.4), 100 μM ONOO^−^ or 100 μM H_2_O_2_ for 1 hour at room temperature in Tris buffer. Fluorescence intensities at 510 nm with a 490 nm excitation were quantified and represented as means ± SD from three independent measurements. Fluorescence enhancements were calculated by dividing the fluorescence of the ONOO^−^- or H_2_O_2_-treated group by that of the control group at the 1-hour time point (DR_ONOO−_ or DR_H2O2_). The selectivity of each mutant was defined as the ratio of the fluorescence enhancement from ONOO^−^-treated group over that from H_2_O_2_-treated group (DR_ONOO−_ / DR_H2O2_). The mutant with the highest selectivity, namely pnGFP1.5-S66A (pnGFP1.6), was chosen for further engineering.

To screen for mutants with further enhanced selectivity, the threonine 5 (Thr5) codon of pnGFP1.6 was mutated into two consecutive degenerative codons (NNKNNK) using primers pnGFP1.6-F-NNK2 and pBAD-RP. The gene library was screened and constructed as described before. The mutant with the highest selectivity from this library was sequenced and designated as pnGFP-Ultra. Characterization protocols for pnGFP-Ultra including the purification, reactivity measurements, selectivity tests over other reactive chemical species, and determination of the limits of detection (LOD) were the same as those for pnGFP (Chen et al., 2013).

### Construction of Mammalian Expression Plasmids

Plasmid *a* and *b* were previously described (Chen et al., 2013). To construct plasmid *c*, the EGFPTAG gene was amplified from plasmid *a* with oligos EGFP-F and EGFP-R. The PCR product was digested with HindIII and ApaI, and ligated into plasmid *b*, which was predigested with the same enzymes. To construct plasmid *d,* the eRF1 gene was first cloned from the total cDNA of HEK 293T cells with oligos eRF1-F and eRF1-R. To prepare the cDNA, total mRNA from HEK293T cell was isolated with the TRI Reagent (Sigma, St. Louis, MO) and reverse-transcribed with the SuperScript^®^ IV Reverse Transcriptase (Thermo Fisher Scientific, Carlsbad, CA). To introduce the E55D point mutation within eRF1, oligos eRF1-NheI-F and E55D-R, E55D-F and eRF1-EcoRI-R were used to amplify the two overlapping fragments flanking E55 position of eRF1, respectively. Next, the two fragments were joined by overlap extension PCR with oligos eRF1-NheI-F and eRF1-EcoRI-R, and sub-cloned into the pUltra vector (Addgene Plasmid #24129) between restriction sites NheI and EcoRI to generate pUltra-eRF1-E55D, which contains the T2A-eRF1-E55D-WPRE cassette. This cassette was amplified with oligos eRF1-GB-F and eRF1-GB-R and ligated into ApaI-linearized plasmid *b* using Gibson assembly cloning (Gibson et al., 2009). To construct plasmid *e*, the CMV-EGFP_TAG_ cassette was amplified from plasmid *a* with oligos CMV-39TAG-XhoI-F and CMV-39TAG-XhoI-R. The PCR product was digested with XhoI and ligated into XhoI digested, DpnI dephosphorylated plasmid *b*. To construct the mammalian expression plasmid for pnGFP-Ultra, the pnGFP-Ultra gene was amplified from pBAD-pnGFP-Ultra with oligos pnGFP-Ultra-HindIII-F and pnGFP-Ultra-ApaI-R, and sub-cloned into plasmid *c* between restriction sites Hind III and ApaI.

### Mammalian Cell Culture, Transfection and Imaging

Human Embryonic Kidney (HEK) 293T cells were cultured in Dulbecco’s Modified Eagle’s Medium (DMEM) supplemented with 10% fetal bovine serum (FBS). Cells were incubated at 37°C with 5% CO2 in humidified air. To test the transfection and non-canonical amino acid (ncAA) incorporation efficiency of the various plasmids, cells were split and seeded into 24-well plates 16 hours before transfection. Cells were transfected with 600 ng DNA and 2 μg PEI (polyethyleneimine, linear, M.W. 25 kDs) per well for three hours, and then cultured in complete medium supplemented with 2 mM DL-*p*BoF for 48 hours. Cells were lysed with 150 μL RIP A buffer and centrifuged at 13,000 rpm, 4°C for 5 mins. The fluorescence of the EGFP-containing supernatant (100 μL) was measured on the microplate reader with excitation and emission wavelength set at 490 nm and 515 nm, respectively. Transfections were triplicated and data were represented as mean ± SD.

To express pnGFP or pnGFP-Ultra, HEK 293T cells in each 35-mm plastic culture dish were co-transfected with 1.5 μg pnGFP-containing plasmid *a* and 1.5 μg plasmid *b* or 1.5 μg pnGFP-Ultra-containing plasmid *c* and 1.5 μg plasmid *d*, in the presence of 10 μg PEI. Other procedures were the same as those for pnGFP (Chen et al., 2013). Cells were imaged under a Leica DMi8 microscope equipped with a Photometrics Prime 95B sCMOS camera. A GFP filter cube with a 470/40 nm bandpass excitation filter and a 525/50 nm bandpass emission filter was used for cell imaging.

### RAW 264.7 Macrophage Cell Culture, Transfection and Activation

RAW 264.7 cells were obtained from ATCC and grown at 37°C under 5% CO_2_ in DMEM supplemented with 10% FBS, 100 μg/mL penicillin, 100 μg/mL streptomycin, and 58 μM β-mercaptoethanol. Cells were passaged 18 h prior to transfection, and next, transfected with 1.5 μg pnGFP-Ultra-containing plasmid *c* and 1.5 μg plasmid *d* in the presence of Viromer according to the manufacturer’s instruction. After transfection, cells were cultured in the aforementioned complete media supplemented with 2 mM DL-*p*BoF. 24 h after transfection, IFN-γ (50 U/ml) and LPS (100 ng/ml) were added into the cell culture media, and the incubation lasted for another 18 h. Cells were next cultured in fresh complete media supplemented with IFN-γ (50 U/ml) and LPS (100 ng/ml) in the absence of DL-*p*BoF for additional 18 h. Cells in the control group were untreated with IFN-γ and LPS but otherwise handled in parallel. Cells were imaged under the same condition as used for imaging HEK 293T.

### Isolation and Culture of Primary Glial Cells

Primary glial cells were isolated from the brain of newborn (0-2 days postnatal) wild-type BALB/C mice and cultured as previously described (Redlich et al., 2013; Seele et al., 2016). Briefly, meninges of mouse brains were removed and whole brains were cut into small pieces and treated with 0.25% trypsin (5 mL/brain) for 15 min at 37°C. After centrifugation (1000 × g, 1 min), trypsin was removed and DMEM (4.5 g/L glucose, 3 mL/brain) was added into the cell precipitation, which was further minced by pipetting up and down with a 1 mL pipette. After another centrifugation (750 × g, 8 min), the supernatant, which contained the needed cells, was filtered with a 40-μm sterile cell strainer (Nylon mesh, Fisherbrand, Cat#: 22363547). Cells were next seeded onto 35 mm cell culture dishes with poly-D-lysine coated glass bottoms and cultured in DMEM (4.5 mg/L glucose) supplemented with 10% FBS, 5 mM glutamine and 100 U/ml penicillin and streptomycin at 37°C, 5% CO_2_ for 8-10 days before transfection. This well-established procedure is known to derive mixed glial cells composed of astrocytes and microglia (Redlich et al., 2013; Seele et al., 2016).

### Transfection and Activation of Primary Glial Cells

Primary glial cells were transfected with Lipofectamine 2000 by following the manufacturer’s instruction. After transfection, cells were cultured in the complete culture media supplemented with 2 mM DL-pBoF. β□amyloid (1-42, human) was added at a final concentration of 5 μM at 24 h post transfection. The incubation lasted for another 24 h and the culture media were replaced with fresh complete media supplemented with 5 μM β□amyloid but without DL-pBoF. The treatment lasted for another 18 h before microscopic images were acquired under the same condition as used for imaging HEK 293T. Cells in the control group were untreated with β□amyloid but otherwise handled in parallel.

## Acknowledgements

Research reported in this publication was supported in part by the National Science Foundation under Award CHE-1750660 and the National Institute of General Medical Science of National Institutes of Health under Awards R01GM118675 and R01GM129291 (including the Supplement Award 3R01GM129291-02S2 from the National Institute on Aging). The content is solely the responsibility of the authors and does not necessarily represent the official views of the funding agencies.

## Author Contributions

H.W., Z.C., and S.Z. conceived and planned experiments; Z.C. and S.Z carried out experiments; X.L isolated and cultured mouse glia cells. Z.C., S.Z., and H.W performed data analysis and interpretation; Z.C. prepared figures and wrote the initial draft of the manuscript; Z.C., S.Z., and H.W. edited and approved the manuscript.

## Supplemental Information

Supplemental Information includes a supplementary table of oligonucleotides sequences used in this study and four supplementary figures and can be found with this article online at http://doi.org/xxx.

