## Supporting Information for "A High-Performance Genetically Encoded Fluorescent Biosensor for Imaging Physiological Peroxynitrite"

Figure S1

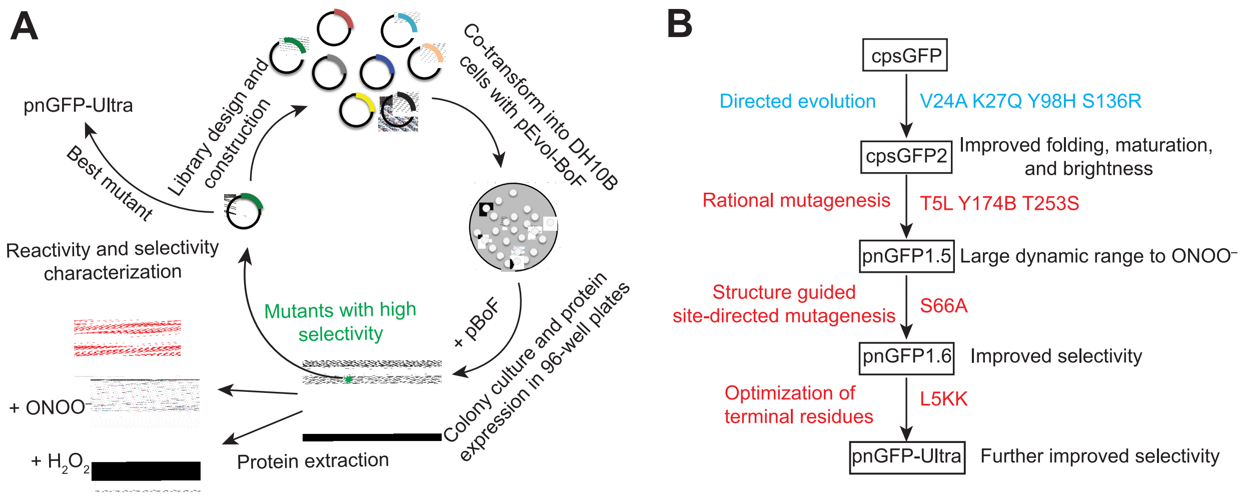


**Figure S1. Screening Procedure and Lineage of pnGFP-Ultra**

(A) Schematic illustration of the screening procedure to derive pnGFP-Ultra. A library of pnGFP mutants were used to co-transformed DH10B electrocompetent cells along with an amber suppression plasmid, pEvol-BoF. Individual colonies were chosen and cultured in 96-well plates with arabinose and *p*BoF to induce protein expression. Protein from individual colonies were extracted and divided into two halves and incubated with $ONOO^-$ or $H_2O_2$ before fluorescent intensities were quantified using a microplate reader. Mutants with high selectivity were selected and used as the template for the next round of screening. After multiple rounds of random and rational mutagenesis, the best mutant pnGFP-Ultra was identified.

(B) Lineage of pnGFP-Ultra. Texts on the left side of the arrow indicate mutagenesis strategy. Texts on the right side of the arrow indicate mutations acquired during the screening process. Texts on the right side of the mutant describe the acquired property of each particular mutant.

Figure S2

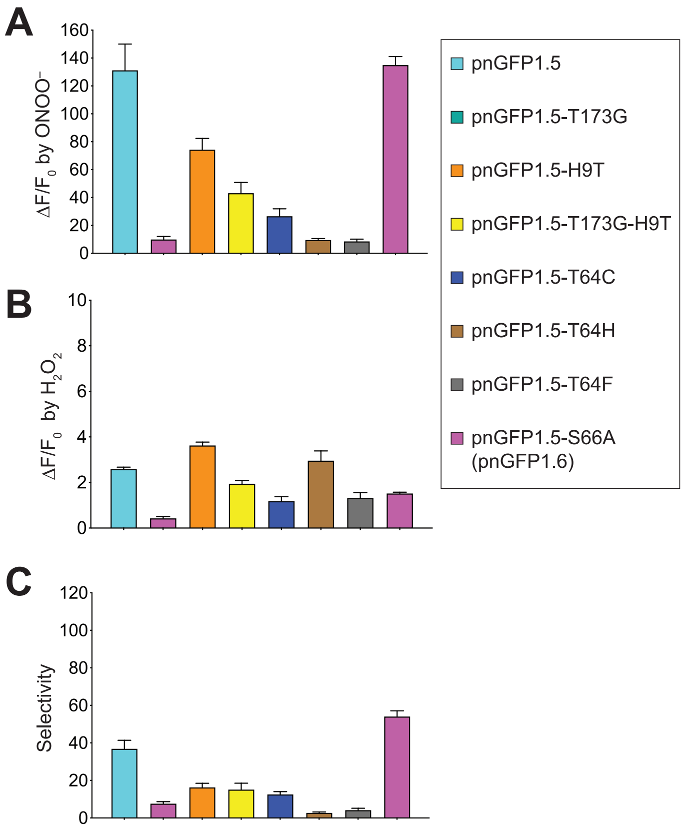


**Figure S2. Characterization of pnGFP1.5 and its mutants**

(A) Dynamic range (expressed as $\Delta F/F_0$) of pnGFP1.5 and relevant mutants towards $ONOO^-$. $F_0$ is the initial fluorescence intensity. $\Delta F$ is the final fluorescent intensity (after treatment) minus $F_0$. Samples were treated with 100 μM $ONOO^-$ for 1 hour. pnGFP1.5, cyan; pnGFP1.5-T173G, green; pnGFP1.5-H9T, orange; pnGFP1.5-T173G-H9T, yellow; pnGFP1.5-T64C, blue; pnGFP1.5-T64H, brown; pnGFP1.5-T64F, grey; pnGFP1.5-S66A, magenta. pnGFP1.5-S66A was later renamed as pnGFP1.6.

(B) Dynamic range ($\Delta F/F_0$) of pnGFP1.5 and relevant mutants towards $H_2O_2$. Samples were treated with 100 μM $H_2O_2$ for 1 hour.

(C) Selectivity of pnGFP1.5 and relevant mutants. Arbitrarily defined selectivity is calculated as the fold of fluorescence enhancement by ONOO− / fold of fluorescence enhancement by $H_2O_2$.

Figure S3

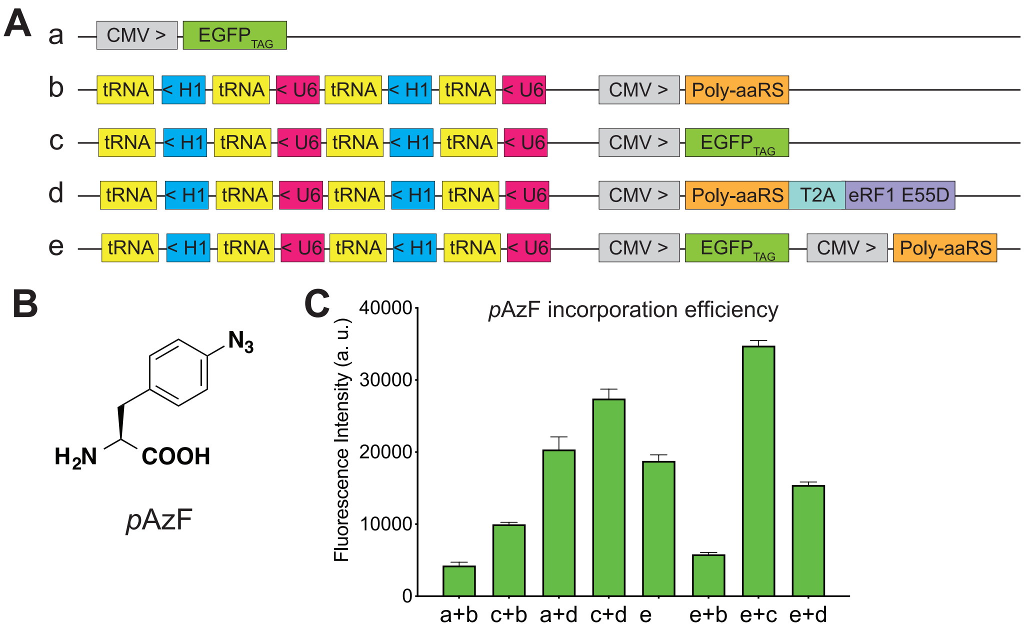


**Figure S3. The Engineered ncAA Incorporation System to also Increase the Incorporation Efficiency of *p*AzF in Mammalian Cells**

(A) Schematics of various expression plasmids. Poly-aaRS is the poly-specific aminoacyl tRNA synthetase, eRF1-E55D is the mutant eRF1 gene, T2A is a self-cleaving 2A peptide, tRNA is the orthogonal tRNA, U6 indicates the U6 promoter, H1 indicates the H1 promoter, CMV is the CMV promoter, $EGFP_{TAG}$ is the EGFP reporter gene with a TAG codon at position 39. > or < denotes the direction of the gene cassettes.

(B) Chemical structure of *p*-azido-phenylalanine (*p*AzF).

(C) Quantification of *p*AzF (1 mM) incorporation into the $EGFP_{TAG}$ reporter gene measured in a cell-based fluorescence assay. The indicated constructs were transiently expressed in HEK 293T cells and the cell lysate green fluorescence was quantified in a plate reader at 515 nm emission with excitation at 490 nm. Data represent the mean ± SD of triplicates.

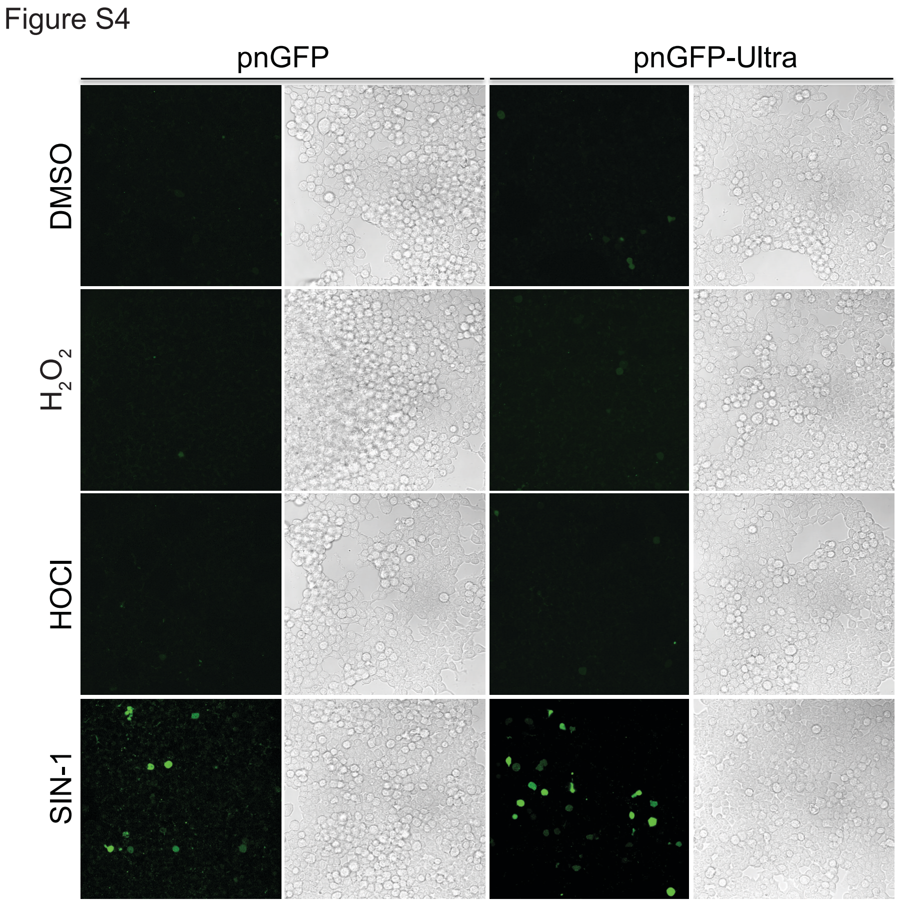


**Figure S4. High Selectivity of pnGFP-Ultra towards Peroxynitrite in Mammalian Cells.** Live-cell imaging of HEK 293T cells transiently expressing pnGFP or pnGFP-Ultra in response to various oxidants. Cells were treated with the indicated chemicals for 90 min before imaging. Concentrations for the chemicals were 1 mM for $H_2O_2$, 100 μM for HOCl, and 100 μM for SIN-1. Images were collected from the fluorescence and bright field channels for each sample.

**Table S1. Oligonucleotides used in this study.**

| Primer name | Nucleotide Sequence |
|---|---|
| pBAD-F | 5'- ATG CCA TAG CAT TTT TAT CC -3' |
| pBAD-R | 5'- GAT TTA ATC TGT ATC AGG -3' |
| pnGFP1.5-T64H-F | 5'- CAC TAC CTG AGC CAC CAG TCC GTG CTG -3' |
| pnGFP1.5-T64H-R | 5'- CAG CAC GGA CTG GTG GCT CAG GTA GTG -3' |
| pnGFP1.5-T64C-F | 5'- CAC TAC CTG AGC TGC CAG TCC GTG CTG -3' |
| pnGFP1.5-T64C-R | 5'- CAG CAC GGA CTG GCA GCT CAG GTA GTG -3' |
| pnGFP1.5-T64F-F | 5'- CAC TAC CTG AGC TTC CAG TCC GTG CTG -3' |
| pnGFP1.5-T64F-R | 5'- CAG CAC GGA CTG GAA GCT CAG GTA GTG -3' |
| pnGFP1.5-S66A-F | 5'- CAC TAC CTG AGC ACC CAG GCC GTG CTG AGC AAA GAC CCC -3' |
| pnGFP1.5-S66A-R | 5'- GGG GTC TTT GCT CAG CAC GGC CTG GGT GCT CAG GTA GTG -3' |
| pnGFP1.5-H9T | 5'- TTT TTG GGC TAA CAG GAG GAA TTA ACC ATG GGC TCG AGC CTG TAC AAC AGC ACC AAG GTC -3' |
| pnGFP1.5-T173G-F | 5'- CTC GTG ACC ACC TTG GGC TAG GGC GTG CAG TGC -3' |
| pnGFP1.5-T173G-R | 5'- GCA CTG CAC GCC CTA GCC CAA GGT GGT CAC GAG -3' |
| pnGFP1.6-F-NNK2 | 5'- TTT TTG GGC TAA CAG GAG GAA TTA ACC ATG GGC TCG AGC NNK NNK TAC AAC AGC CAC AAG -3' |
| EGFP-F | 5'- TAC TGA AGC TTG CCG CCA CCA TGG TGA GCA AGG GCG AGG AGC TG -3' |
| EGFP-R | 5'- TAC TGA GGG CCC TTA TTA CTT GTA CAG CTC GTC CAT GCC -3' |
| eRF1-F | 5'- ATG GCG GAC GAC CCC AGT GCT GCC GAC AGG AAC GTG GAG ATC TGG AAG ATC AAG AAG CTC -3' |
| eRF1-R | 5'- CTA GTA GTC ATC AAG GTC AAA AAA TTC ATC GTC TCC TCC TTG GTA TTC CAT TCC -3' |
| E55D-F | 5'- AAT GTT AGC GGA TGA TTT TGG AAC TGC AT -3' |
| E55D-R | 5'- ATG CAG TTC CAA AAT CAT CCG CTA ACA TT -3' |
| eRF1-NheI-F | 5'- ATA CGC TAG CAT GGC GGA CGA CCC CAG TGC TGC CGA CAG GAA CGT G -3' |
| eRF1-EcoRI-R | 5'- ATA GTG AAT TCT CAT TAG TAG TCA TCA AGG TCA AAA AAT TCA TCG TCT CCT CC -3' |
| eRF1-GB-F | 5'- TAC TGT CTG ATT TGC TGG AAA GGG CCC GTT GGA TCC GGC TCC GGC GAG GGC AGG GGA AGT -3' |
| eRF1-GB-R | 5'- CAC AGT CGA GGC TGA TCA GCG GGT TTA AAC GCG GGG AGG CGG CCC AAA GGG AGA TCC GAC -3' |

| | |
|---|---|
| CMV-39TAG-XhoI-F | 5'- TGT TGG CTC GAG TCT AGA TAT ACG CGT TGA CAT TGA TTA T -3' |
| CMV-39TAG-XhoI-R | 5'- TCA ATG CTC GAG AAG CCA TAG AGC CCA CCG CAT CCC CAG -3' |
| PnGFP-Ultra-HindIII-F | 5'- TAT ACA AGC TTG CCG CCA CCA TGG GCT CCA GCA AGA AGT ACA ACA GC -3' |
| PnGFP-Ultra-ApaI-R | 5'- GAA TAG GGC CCT TAT TAA TGG TGA TGG TGA TGG TG -3' |
